# Soil microbes mediate the effects of nitrogen supply and co-inoculation on Barley Yellow Dwarf Virus in *Avena sativa*

**DOI:** 10.1101/2021.04.28.441777

**Authors:** Casey A. Easterday, Amy E. Kendig, Christelle Lacroix, Eric W. Seabloom, Elizabeth T. Borer

## Abstract

- Nutrient supply rates to hosts can mediate host–pathogen interactions. In terrestrial systems, nutrient supply to plants is mediated by soil microbes, suggesting a potential indirect effect of soil microbes on plant–pathogen interactions. Soil microbes also may affect plant pathogens by inducing plant defenses.
- We tested the role of soil microbes, nitrogen supply to plant hosts, and co-inoculation on infection by aphid-vectored RNA viruses, Barley Yellow Dwarf Virus (BYDV-PAV) and Cereal Yellow Dwarf Virus (CYDV-RPV), in a grass host grown in soil microbes collected from a long-term nitrogen enrichment experiment.
- BYDV-PAV incidence declined with high nitrogen supply, co-inoculation, or presence of soil microbes exposed to long-term low nitrogen enrichment. However, when combined, the negative effects of these treatments were sub-additive: nitrogen and co-inoculation did not reduce BYDV-PAV incidence in plants grown with the soil microbes. While soil microbes impacted leaf chlorophyll, they did not alter biomass or CYDV-RPV incidence.
- Soil microbes mediated the effects of nitrogen supply and co-inoculation on infection incidence and the effects of infection on host symptoms. Thus, soil microbial communities can indirectly control disease dynamics, altering the effects of nitrogen enrichment on plant–pathogen and pathogen–pathogen interactions in terrestrial systems.

## Introduction

Fossil fuel use and fertilizer production have more than doubled reactive nitrogen (N) inputs to terrestrial ecosystems since pre-industrialization (Vitousek *et al*., 1997; Galloway *et al*., 2004). Nitrogen enrichment can profoundly impact terrestrial plant systems, increasing productivity and reducing biodiversity (Elser *et al*., 2007; Midolo *et al*., 2019). Further, the effects of N enrichment on plants can have repercussions throughout food webs (Sedlacek *et al*., 1988; Ritchie, 2000; He & Silliman, 2015). For example, N enrichment can modify plant– pathogen interactions (Dordas, 2009; Veresoglou *et al*., 2013) and interactions among different pathogens that co-infect plants (Lacroix *et al*., 2014; Kendig *et al*., 2020). The communities of pathogens that rely on plants can in turn impact plant productivity, community composition, and ecosystem processes (Lovett *et al*., 2010; Paseka *et al*., 2020; Borer *et al*., 2021). However, a key component of terrestrial systems—soil microbes—have been neglected in many studies of N enrichment on aboveground plant pathogens. Soil microbes affect plant access to N (van der Heijden *et al*., 2008; Kuzyakov & Xu, 2013) and plant interactions with above- and belowground pathogens (van Loon *et al*., 1998; Berendsen *et al*., 2012). Therefore, laboratory-based studies that do not include natural soil microbial communities may under- or over-estimate the effects of N enrichment on plant–pathogen interactions under field conditions. Moreover, N enrichment can change the nature of interactions between soil microbes and plants through time (Johnson, 1993; Keeler *et al*., 2008; Weese *et al*., 2015; Huang *et al*., 2019). Consequently, even studies that include natural soil microbial communities may mis-characterize the impacts of N enrichment on plant pathogen communities if they do not encompass long enough time scales.

Nitrogen is an essential component of the genetic material and proteins in plants and microbes (Sterner & Elser, 2002). Changes in N availability can therefore modify the fitness of plants (Johnson, 1993; Welch & Leggett, 1997), microbes (Schimel & Bennett, 2004; Kuzyakov & Xu, 2013), and insect vectors of plant pathogens (Nowak & Komor, 2010; Bogaert *et al*., 2017). Because plants, their pathogens, and the insect vectors of pathogens rely on N, N enrichment can increase or decrease infection prevalence (Seabloom *et al*., 2010; Borer *et al*., 2014), pathogen load (Singh, 1970; Hoffland *et al*., 2000; Mitchell *et al*., 2003; Robert *et al*., 2004; Fagard *et al*., 2014; Whitaker *et al*., 2015) and disease resistance (Dietrich *et al*., 2004; Bellin *et al*., 2013; Mur *et al*., 2016). Further, individual plants and plant communities frequently host multi-pathogen communities (Seabloom *et al*., 2009; Bass *et al*., 2019) which may shift in composition with N enrichment (Lacroix *et al*., 2014; Kendig *et al*., 2020). Soil microbes and N enrichment may interact to affect plant–pathogen and pathogen–pathogen interactions. For example, fertilizer and green manure can increase the disease-suppressive activity of foliar and rhizosphere microbial communities (Wiggins & Kinkel, 2005; Berg & Koskella, 2018).

Soil microbes may mediate plant–pathogen interactions by altering the amount of N available to the plant. Soil microbes can increase plant access to N through N fixation, N mineralization, and extending root networks (van der Heijden *et al*., 2008), and they also may compete with plants for N (Schimel & Bennett, 2004; Kuzyakov & Xu, 2013). N enrichment can reduce the benefits plants receive from microbial mutualists (Johnson, 1993; Weese *et al*., 2015), although losses can be offset by the direct benefits of bioavailable N (Johnson, 1993; Farrer & Suding, 2016). N enrichment has variable effects on N mineralization rates (Mueller *et al*., 2013; Chen *et al*., 2019), mediated by changes in soil microbial community composition and pH (Chen *et al*., 2019). Over decadal time scales, N enrichment can drive large compositional and evolutionary changes in soil microbial communities that affect their N-related interactions with plants (Klinger *et al*., 2016; Huang *et al*., 2019). For example, N enrichment can shift the relative abundance of archaea and bacteria that oxidize ammonia, depending on the form of N added (Leff *et al*., 2015; Moreau *et al*., 2015).

Soil microbial communities can contain plant pathogens as well as microbes that suppress plant pathogens (Schlatter *et al*., 2017). Soil microbes can suppress soil-borne pathogens via competition for resources, interference with pathogen signaling, or production of antibiotic compounds and lytic enzymes (Lugtenberg & Kamilova, 2009; Berendsen *et al*., 2012). In addition, beneficial and pathogenic soil microbes can induce disease resistance pathways in plants, priming them for faster and stronger responses to aboveground pathogen attacks (van Loon *et al*., 1998; Pieterse *et al*., 2014; Mauch-Mani *et al*., 2017). Soil biota associated with induced disease resistance include bacteria in the genera *Pseudomonas, Serratia*, and *Bacillus* and fungi in the genera *Trichoderma, Fusarium, Piriformospora*, and *Glomeromycota* (Pieterse *et al*., 2014; Mauch-Mani *et al*., 2017). Nitrogen enrichment may modify the effects of soil microbes on plant diseases through compositional or evolutionary shifts in microbial communities or changes in microbe–pathogen interactions (Otto-Hanson *et al*., 2013; Schlatter *et al*., 2013; Klinger *et al*., 2016; Huang *et al*., 2019). For example, N enrichment can reduce the abundance and colonization rates of arbuscular mycorrhizal fungi (AMF), which includes the genus *Glomeromycota* (Treseder, 2004; Leff *et al*., 2015; Jia *et al*., 2020). In contrast, N enrichment tends to increase phyla and classes containing some groups of fungi (*Trichoderma* and *Fusarium*) and bacteria (*Pseudomonas, Serratia*, and *Bacillus*) that induce disease resistance (Fierer *et al*., 2012; Ramirez *et al*., 2012; Leff *et al*., 2015; Chen *et al*., 2019).

Here, we evaluated the effects of soil microbial communities on N-dependent changes in plant–pathogen and pathogen–pathogen interactions, using two widespread and economically important insect-vectored plant viruses (BYDV-PAV and CYDV-RPV). We used aphids (*Rhopalosiphum padi*) to inoculate oat plants (*Avena sativa*) with these Barley and Cereal Yellow Dwarf Viruses (B/CYDV’s) across a full factorial combination of N supply rates (low or high) and soil microbial communities collected from soils subjected to different levels of long-term N fertilization rates (ambient, low, or high). For each treatment, we measured virus incidence (i.e., the proportion of plants that became infected), plant biomass, and leaf chlorophyll content. We addressed three questions: (1) What are the effects of N on single infection and co-infection incidence in plants grown in sterile soil? (2) Do soil microbes mediate the effects of N on single infection and co-infection incidence (3) Do soil microbes mediate the effects of N or infection on host traits?

## Materials and Methods

### Study system

B/CYDVs, generalist pathogens of the Luteoviridae family, cause systemic infections in over 150 grass species in the Poacaea family, stunting growth, yellowing or reddening leaves, and reducing fecundity (Carrigan *et al*., 1983; Irwin & Thresh, 1990; D’Arcy & Burnett, 1995). B/CYDVs comprise members of the Luteovirus (BYDVs) and Polerovirus (CYDVs) genera (Miller *et al*., 2002). BYDV-PAV and CYDV-RPV are considered representative members of each genera and have been the foci of many studies (Power *et al*., 1991; Seabloom *et al*., 2009; Lacroix *et al*., 2014; Kendig *et al*., 2020). These viruses are transmitted by a range of aphid vectors, including *Rhopalosiphum padi* that transmits both BYDV-PAV and CYDV-RPV (D’Arcy & Burnett, 1995). Importantly, these viruses are strictly insect-vectored; they cannot be transmitted to plants via soil. We used *A. sativa* L. cv. Coast Black Oat as our plant host species. *Avena sativa* can host AMF (Yang *et al*., 2010) and AMF were found to have a positive effect on BYDV-PAV titer in *Avena fatua* under elevated CO_2_ (Rúa *et al*., 2013). Soil microbes were collected from Cedar Creek Ecosystem Science Reserve and Long-Term Ecological Research (LTER) site (CDR; see next section). Prior to conducting this work, we established that both of our focal viruses are present in natural communities at CDR. In 2009, we haphazardly collected 153 plants of 10 different grass species across CDR to determine which B/CYDV’s were present. We assessed infection status via ELISA as detailed in Seabloom et al. (2009). Overall infection prevalence of PAV was 0.17 and RPV was 0.03.

### Soil microbes

In June 2014, we collected soil cores from a long-term experiment in a successional grassland at CDR (experiment “E001”, www.cedarcreek.umn.edu; Bethel, MN, USA). Cedar Creek has sandy, N-limited soils and a background wet N deposition rate of approximately 6 kg N ha^-1^ year^-1^ (58% NH_4_, 42% NO_3_) (Tilman, 1987; Clark & Tilman, 2008). We collected soils from field A, which was abandoned from agriculture in 1968 and burned annually beginning in 2005. These plots had received annual additions of P, K, Ca, Mg, S, and citrate-chelated trace metals since 1982 and three levels of N fertilizer: 0, 34, or 270 kg N ha^-1^ yr^-1^ (see Tilman, 1987 for details). In this experiment, N fertilization has increased plant biomass and soil N concentration and decreased plant species richness (Isbell *et al*., 2013a,b). We randomly sampled three plots for each N fertilization rate (Fig. **S1**) and six locations within each 4 x 4 m plot to extract a soil core (1.9 cm diameter and 10 cm deep). In the lab, soil cores were passed through a 4 mm sieve, then twice through a 2 mm sieve to remove coarse debris and roots, and then combined based on their N fertilization rate.

Next, we prepared soil microcosms by filling four large, surface sterilized bins with 17 L of potting soil composed of 70% Sunshine medium vermiculite (vermiculite and less than 1% crystalline silica; Sun Gro Horticulture, Agawam, MA, USA) and 30% Turface MVP (calcined clay containing up to 30% crystalline silica; Turface Athletics, Buffalo Grove, IL, USA), saturated with tap water (approximately 5 L for every 20 L of dry soil) and autoclaved at 121°C and 15 psi for 60 minutes to kill the naturally existing microbial consortium. We then mixed 350 mL of field soil from each N fertilization level separately into the bins. Field soil comprised approximately 2% of the bin soil volume. We did not mix field soil into the fourth bin. Lastly, we covered the bins with non-airtight lids and incubated the soil at 25°C for 11 days.

### Experimental setup and implementation

For each of the four soil microcosms, we filled 80 conical plastic pots (3.8 cm diameter x 21 cm depth, 164 ml) with soil mixture and planted one *A. sativa* seed per pot 4.5 cm from the surface of the soil. Seeds were obtained from the USDA (National plant germplasm system, USDA; USA) in June 2013 and were surface sterilized with 12.5% bleach solution. Then, we haphazardly assigned plants to later receive one of two N supply rates (7.5 μM NH_4_NO_3_ was “low N” and 375 μM NH_4_NO_3_ was “high N”; Table **S1**) and one of four virus inoculations (BYDV-PAV, CYDV-RPV, co-inoculation, or mock inoculation), leading to ten replicates per treatment. Plants grew in a growth chamber containing only healthy plants with a 16:8 h light:dark cycle at 19-20°C under Lumilux high pressure sodium ET18 bulbs for 11 days. Two days after planting, we watered the pots with 30 ml of the modified Hoagland solution (Hoagland & Arnon, 1938; Lacroix *et al*., 2014; Table **S1**) corresponding to the plant’s assigned N supply rate. We watered plants with these solutions twice per week until harvest.

When the plants had been growing for 22 days, we used *R. padi* aphids to inoculate them with BYDV-PAV, CYDV-RPV, both viruses, or to perform a mock inoculation. *Rhopalosiphum padi* were obtained from Dr. G. Heimpel at the University of Minnesota (St. Paul, MN, USA) and reared on *A. sativa* in growth chamber conditions described above (except with 28W Ultramax EcoXL lights). BYDV-PAV and CYDV-RPV isolates were obtained from Dr. S. Gray at Cornell University (Ithaca, NY, USA) in January 2013. They were also maintained in *A. sativa* plants in similar growth chamber conditions (except with 40W cool white light bulbs). We inoculated plants by allowing aphids to feed on either BYDV-PAV- or CYDV-RPV-infected *A. sativa* tissue in 25 mL glass tubes sealed with corks for approximately 48 hours. Then, we transferred the aphids to 2.5 x 8.5 cm, 118 μm polyester mesh cages secured to one leaf on each experimental plant with Parafilm and bobby pins. Ten aphids were used to inoculate each plant, with 5 carrying each virus for the co-inoculation treatment, five viruliferous (carrying virus) and five non-viruliferous aphids for each single virus treatment, and ten non-viruliferous aphids for the mock inoculation treatment. We allowed aphids to feed on the experimental plants for approximately 96 hours, after which we manually killed all aphids and removed the cages. Plants grew for 19 more days before we took measurements. To estimate N stress through leaf chlorophyll content (Zhao *et al*., 2015), we took three measurements per plant with a SPAD-502 Meter (Soil Plant Analysis Development; Konica Minolta, Tokyo, Japan). Then, we harvested and weighed the aboveground biomass, which we stored at −20°C until it was analyzed for virus infection.

### Detection of B/CYDV infection

To extract total RNA, we ground approximately 50 mg of leaf tissue in a bead-beater with a copper BB and 1 ml of TRIzol™ Reagent (Invitrogen™, Thermo Fisher Scientific, Waltham, MA, USA) per the manufacturer’s instructions. We then purified RNA from the cellular components following the extraction protocol published by Lacroix et. al. (2014). We re-suspended the purified RNA in nuclease-free water and stored the samples at −20°C until performing the reverse transcription polymerase chain reaction (RT-PCR). We used a nanodrop spectrophotometer (Thermo Fisher Scientific) to quantify the concentration of RNA within each sample and then performed a multiplex RT-PCR assay to isolate and amplify BYDV-PAV and CYDV-RPV nucleic acids as published previously (Deb & Anderson, 2008; Lacroix *et al*., 2014).

We combined 5 μl of each PCR product with 2 μl of 6X loading dye (Genesee Scientific, El Cajon, CA, USA) and loaded the samples and 100 bp DNA ladder (Apex Bioresearch Products, North Liberty, IA, USA) into an Agarose-1000 gel (Invitrogen, Thermo Fisher Scientific) stained with 2% SybrSafe (Invitrogen, Thermo Fisher Scientific). After 25 minutes at 120 V, we observed the gel with a UV-light EZ doc system (Bio-Rad Laboratories, Hercules, CA, USA) to detect bands at 298 bp and 447 bp, indicating the presence of BYDV-PAV and CYDV-RPV, respectively.

### Statistical analyses

We assessed the effects of the experimental treatments on the infection incidence of BYDV-PAV and CYDV-RPV (i.e., the proportion of plants infected out of those inoculated) using binomial (logit-link) generalized linear regressions with virus infection as a binary response variable and soil microbe inoculum (sterilized, ambient N, low N, or high N), N supply (binary variable), whether the plants were co-inoculated (binary variable), and their interactions as independent variables. The intercepts represented singly inoculated plants grown in sterile soil with low N supply. We tested the effects of N supply and soil microbe inoculum on co-infection incidence using an analogous procedure. Samples with an infection inconsistent with the inoculation treatment were removed from analyses. Inconsistent infections likely arose from small aphids escaping cages during the inoculation period and occurred in 31 of 229 plants (Table **S2**). Treatment sample sizes in the final dataset ranged from seven to ten.

To assess the effects of the experimental treatments on the *A. sativa* plants, we used linear regressions with log-transformed biomass and log-transformed chlorophyll content as response variables and N supply, soil microbe inoculum, successful inoculation treatment (mock, BYDV-PAV only, and CYDV-RPV only), and their interactions as the independent variables. Therefore, we omitted plants from analyses that were unsuccessfully inoculated, either because the intended infection was not detected or because an unintended infection was detected (Table **S2**). Co-infected plants were omitted from analyses due to limited sample sizes. The chlorophyll values used in the model were the averages of three measurements taken per plant. The intercepts represented mock-inoculated plants grown in sterile soil with low N supply. Treatment sample sizes in the final dataset ranged from three to nine.

All regressions described above were fit using Bayesian models with the brms package in R version 4.0.2 (Bürkner, 2017; R Core Team, 2020). Models had three chains of 6000 iterations each with a 1000 iteration burn-in period. Gaussian distributions with a mean of zero and a standard deviation of ten were used as prior distributions for intercepts and coefficients (very weakly informative; McElreath, 2015). We used a half Student’s *t*-distributions with three degrees of freedom, a location of zero, and a scale of ten as the prior distribution for the residual standard deviations (Bürkner, 2017). We assessed model fit by ensuring that r-hat values were equal to one, that the three chains were well mixed, and that simulated data from the posterior predictive distributions were consistent with observed data. In the results, we present point estimates with 95% highest posterior density intervals based on posterior samples of model coefficients in brackets.

To evaluate the effect of sample size on the probability of detecting an effect with quantile-based 95% credible intervals that omit zero, we simulated 1000 datasets of the same sample sizes and with the mean effect size measured in the experiment. We fit regressions to each dataset and calculated the number of times the 95% credible intervals of the variable of interest omitted zero (Kurz, 2019). We repeated the analysis with multiple sample sizes. We performed this analysis for the effects of CYDV-RPV infection on log-transformed plant biomass, where the mean difference was −0.23, the sample sizes were 8 (mock-inoculated, low N supply, sterile soil) and 6 (CYDV-RPV infected, low N supply, sterile soil), and the regression was a normal linear regression with infection status as the independent variable.

## Results

### The effects of N on infection incidence in sterile soils

BYDV-PAV incidence of singly inoculated plants grown with low N supply was 0.96 [0.84, 1.00] (Fig. **1a**). High N supply reduced BYDV-PAV incidence in singly inoculated plants to 0.61 (−36% [-72%, −4.0%]) and co-inoculation reduced BYDV-PAV incidence to 0.32 (−66% [-93%, −38%], Fig. **1a**). However, high N supply did not affect BYDV-PAV incidence of co-inoculated plants (estimated change relative to low N supply: 56% [-83%, 283%]), leading to an interaction between N supply and co-inoculation (Table **1**). CYDV-RPV incidence of singly inoculated plants grown with low N supply was 0.66 [0.38, 0.93] (Fig. **2a**). Co-inoculation reduced CYDV-RPV incidence to 0.11 (−83% [-100%, −53%], Fig. **2a**). Nitrogen supply did not affect CYDV-RPV incidence in singly or co-inoculated plants (Table **2**). The average co-infection incidence of co-inoculated plants grown in sterile soil with low N supply was 0.10 [0.00, 0.28]. Nitrogen supply did not affect co-infection incidence (Fig. **3a**, Table **3**).

**Table 1.** Generalized linear model summary of BYDV-PAV incidence (*n* = 139).

| Variable<sup>a</sup> | Estimate | Std. Error | 95% CI |  | Rhat |
| --- | --- | --- | --- | --- | --- |
|  |  |  | Lower | Upper |  |
| **intercept<sup>b</sup>** | **4.12** | **1.73** | **1.36** | **8.04** | **1.00** |
| **N supply** | **-3.60** | **1.85** | **-7.66** | **-0.43** | **1.00** |
| **co-inoculation** | **-4.95** | **1.82** | **-8.97** | **-1.92** | **1.00** |
| soil ambient N | -2.96 | 1.90 | -7.09 | 0.43 | 1.00 |
| **soil low N** | **-4.27** | **1.84** | **-8.37** | **-1.20** | **1.00** |
| soil high N | 6.48 | 4.99 | -1.67 | 17.51 | 1.00 |
| **N supply x co-inoculation** | **3.92** | **2.01** | **0.34** | **8.19** | **1.00** |
| soil ambient N x N supply | 2.09 | 2.14 | -1.93 | 6.48 | 1.00 |
| **soil low N x N supply** | **5.84** | **2.24** | **1.88** | **10.64** | **1.00** |
| soil high N x N supply | 3.53 | 5.62 | -6.95 | 15.09 | 1.00 |
| **soil ambient N x co-inoculation** | **3.77** | **2.06** | **0.02** | **8.13** | **1.00** |
| **soil low N x co-inoculation** | **5.05** | **2.02** | **1.43** | **9.40** | **1.00** |
| soil high N x co-inoculation | -4.70 | 5.02 | -15.72 | 3.66 | 1.00 |
| soil ambient N x N supply x co-inoculation | -0.78 | 2.50 | -5.87 | 3.95 | 1.00 |
| soil low N x N supply x co-inoculation | -4.48 | 2.59 | -9.75 | 0.42 | 1.00 |
| soil high N x N supply x co-inoculation | -3.34 | 5.68 | -15.05 | 7.19 | 1.00 |

<sup>a</sup>Variables with quantile-based 95% credible intervals (CI) that omit zero are in bold.
<sup>b</sup>Intercept factor levels: low N supply, single inoculation, sterile soil

**Table 2.**
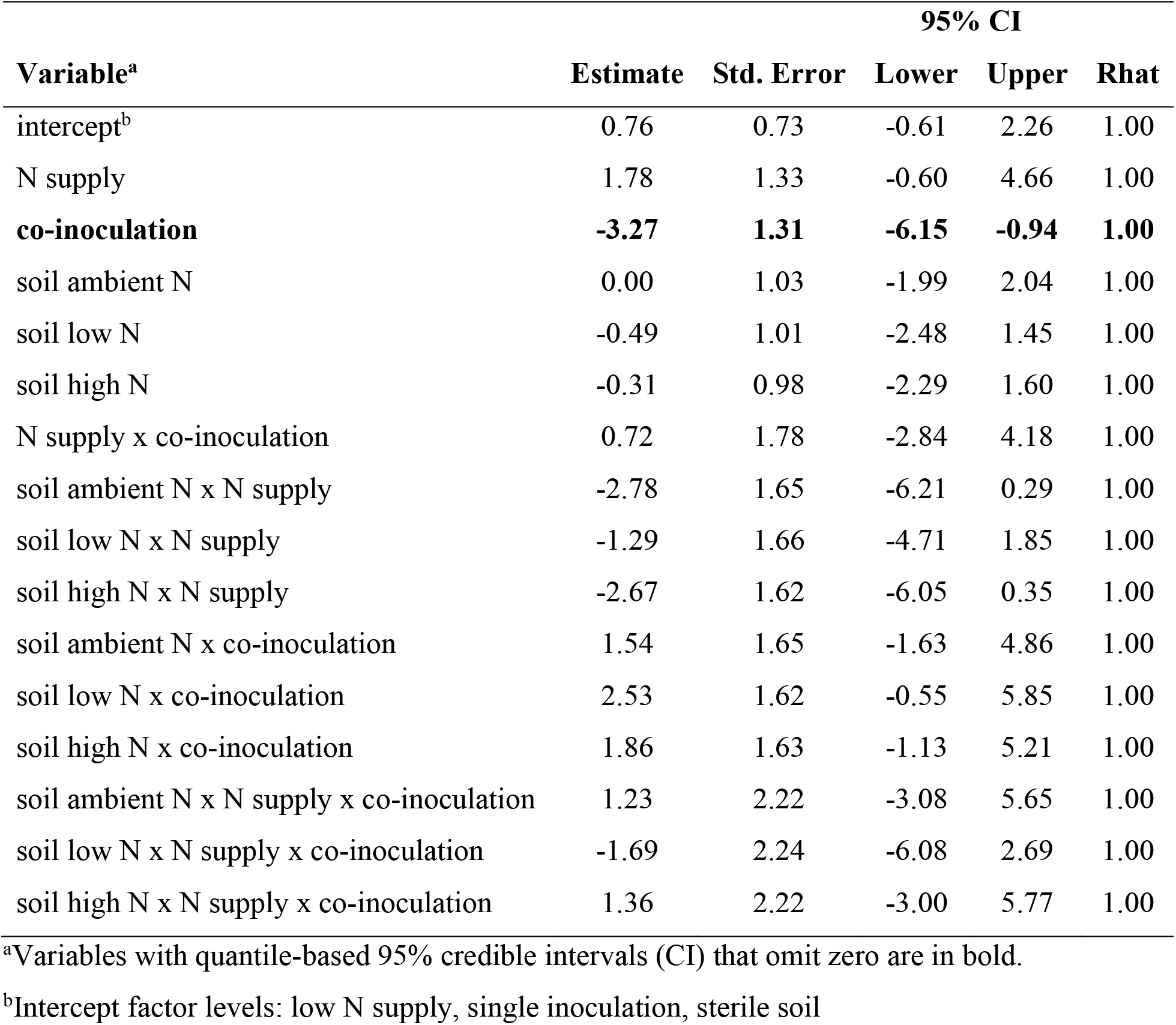
Generalized linear model summary of CYDV-RPV incidence (*n* = 154).

| Variable<sup>a</sup> | Estimate | Std. Error | 95% CI |  | Rhat |
| --- | --- | --- | --- | --- | --- |
|  |  |  | Lower | Upper |  |
| intercept<sup>b</sup> | 0.76 | 0.73 | -0.61 | 2.26 | 1.00 |
| N supply | 1.78 | 1.33 | -0.60 | 4.66 | 1.00 |
| **co-inoculation** | **-3.27** | **1.31** | **-6.15** | **-0.94** | **1.00** |
| soil ambient N | 0.00 | 1.03 | -1.99 | 2.04 | 1.00 |
| soil low N | -0.49 | 1.01 | -2.48 | 1.45 | 1.00 |
| soil high N | -0.31 | 0.98 | -2.29 | 1.60 | 1.00 |
| N supply x co-inoculation | 0.72 | 1.78 | -2.84 | 4.18 | 1.00 |
| soil ambient N x N supply | -2.78 | 1.65 | -6.21 | 0.29 | 1.00 |
| soil low N x N supply | -1.29 | 1.66 | -4.71 | 1.85 | 1.00 |
| soil high N x N supply | -2.67 | 1.62 | -6.05 | 0.35 | 1.00 |
| soil ambient N x co-inoculation | 1.54 | 1.65 | -1.63 | 4.86 | 1.00 |
| soil low N x co-inoculation | 2.53 | 1.62 | -0.55 | 5.85 | 1.00 |
| soil high N x co-inoculation | 1.86 | 1.63 | -1.13 | 5.21 | 1.00 |
| soil ambient N x N supply x co-inoculation | 1.23 | 2.22 | -3.08 | 5.65 | 1.00 |
| soil low N x N supply x co-inoculation | -1.69 | 2.24 | -6.08 | 2.69 | 1.00 |
| soil high N x N supply x co-inoculation | 1.36 | 2.22 | -3.00 | 5.77 | 1.00 |

<sup>a</sup>Variables with quantile-based 95% credible intervals (CI) that omit zero are in bold.
<sup>b</sup>Intercept factor levels: low N supply, single inoculation, sterile soil

**Figure 1.**
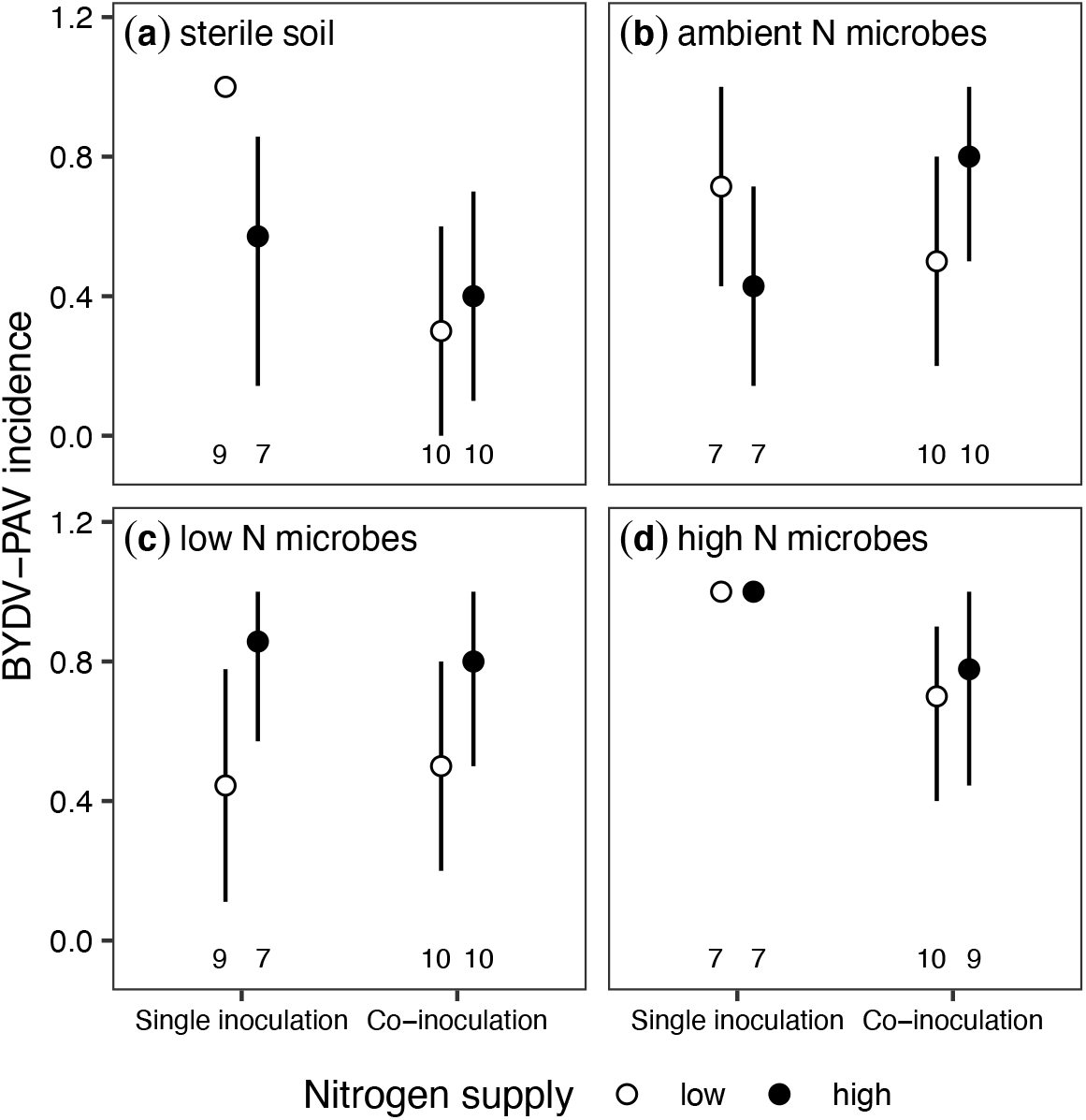
BYDV-PAV incidence (mean ± 95% confidence intervals) of plants grown with (a) sterile soil, (b) microbes exposed to long-term ambient N, (c) microbes exposed to long-term low N, and (d) microbes exposed to long-term high N. Plants were grown with low or high N supply and were either singly- or co-inoculated. Corresponding sample sizes are labelled under points and error bars.

**Figure 2.**
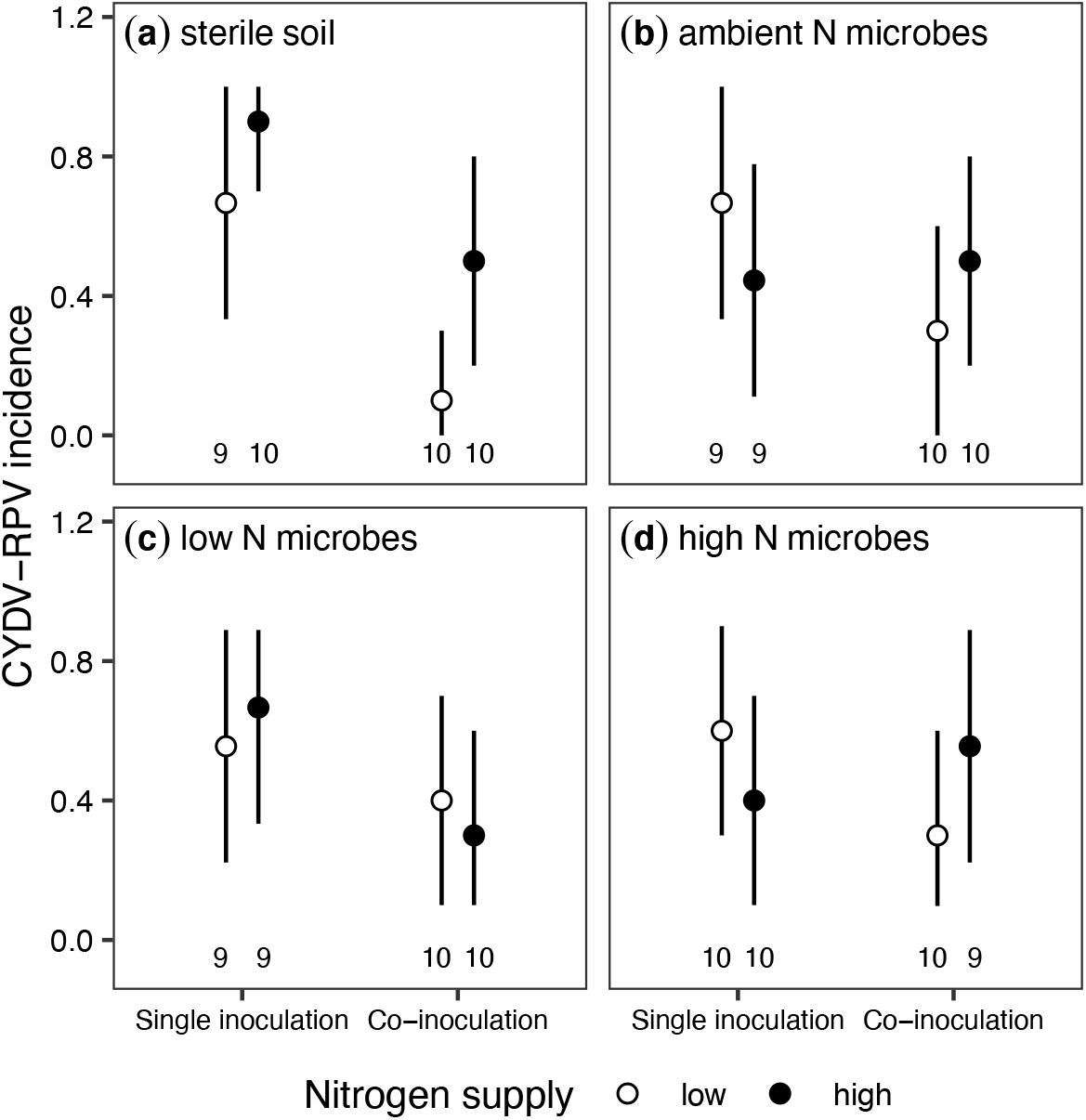
CYDV-RPV incidence (mean ± 95% confidence intervals) of plants grown with (a) sterile soil, (b) microbes exposed to long-term ambient N, (c) microbes exposed to long-term low N, and (d) microbes exposed to long-term high N. Plants were grown with low or high N supply and were either singly- or co-inoculated. Corresponding sample sizes are labelled under points and error bars.

### Soil microbes x N: effects on infection incidence

Inoculation with microbes from low N fertilization field soils (“low N microbes”) reduced BYDV-PAV incidence to 0.47 [0.18, 0.77], a 51% decrease [-83%, −20%] relative to singly inoculated plants grown in sterile soils with low N supply (Fig. **1c**). In contrast to the negative effect of high N supply when plants were grown in sterile soil (Fig. **1a**), high N did not affect BYDV-PAV incidence when plants were grown with low N microbes (estimated change relative to low N supply: 109% [-37%, 310%], Fig. **1c**), leading to an interaction between N supply and low N microbes (Table **1**). Similarly, high N supply did not affect BYDV-PAV incidence when plants were grown with ambient N microbes (estimated change relative to low N supply: −38% [-91%, 22%], Fig. **1b**) or high N microbes (estimated change relative to low N supply: −0.27% [-9.4%, 5.4%], Fig. **1d**), but there was no statistical interaction between N supply and these microbe treatments (Table **1**). In contrast to plants grown with sterile soil (Fig. **1a**), co-inoculation did not affect BYDV-PAV incidence when plants were grown with low N microbes (estimated change relative to single inoculation: 21% [-75%, 158%], Fig. **1c**) or ambient N microbes (estimated change relative to single inoculation: −27% [-80%, 32%], Fig. **1b**), leading to interactions between co-inoculation and each of the microbe treatments (Table **1**). However, co-inoculation reduced BYDV-PAV incidence from 0.99 to 0.70 (−29% [-57%, −4.9%]) when plants were grown with high N microbes (Fig. **1d**). High N supply did not affect BYDV-PAV incidence of co-inoculated plants grown with any of the microbe treatments (Table **1**). Microbes in field soils did not affect CYDV-RPV incidence (Fig. **2b–d**, Table **2**) or co-infection incidence (Fig. **3b–d**, Table **3**) relative to sterile soil.

**Table 3.** Generalized linear model summary of co-infection incidence (*n* = 79).

| Variable<sup>a</sup> | Estimate | Std. Error | 95% CI |  | Rhat |
| --- | --- | --- | --- | --- | --- |
|  |  |  | Lower | Upper |  |
| **intercept<sup>b</sup>** | **-2.60** | **1.16** | **-5.23** | **-0.72** | **1.00** |
| N supply | 1.01 | 1.42 | -1.67 | 3.94 | 1.00 |
| soil ambient N | -0.07 | 1.70 | -3.58 | 3.25 | 1.00 |
| soil low N | 0.99 | 1.45 | -1.70 | 4.04 | 1.00 |
| soil high N | 0.99 | 1.44 | -1.70 | 4.01 | 1.00 |
| soil ambient N x N supply | 1.65 | 1.99 | -2.12 | 5.71 | 1.00 |
| soil low N x N supply | -0.35 | 1.81 | -4.01 | 3.12 | 1.00 |
| soil high N x N supply | 0.34 | 1.79 | -3.24 | 3.73 | 1.00 |

<sup>a</sup>Variables with quantile-based 95% credible intervals (CI) that omit zero are in bold.
<sup>b</sup>Intercept factor levels: low N supply, sterile soil

**Figure 3.**
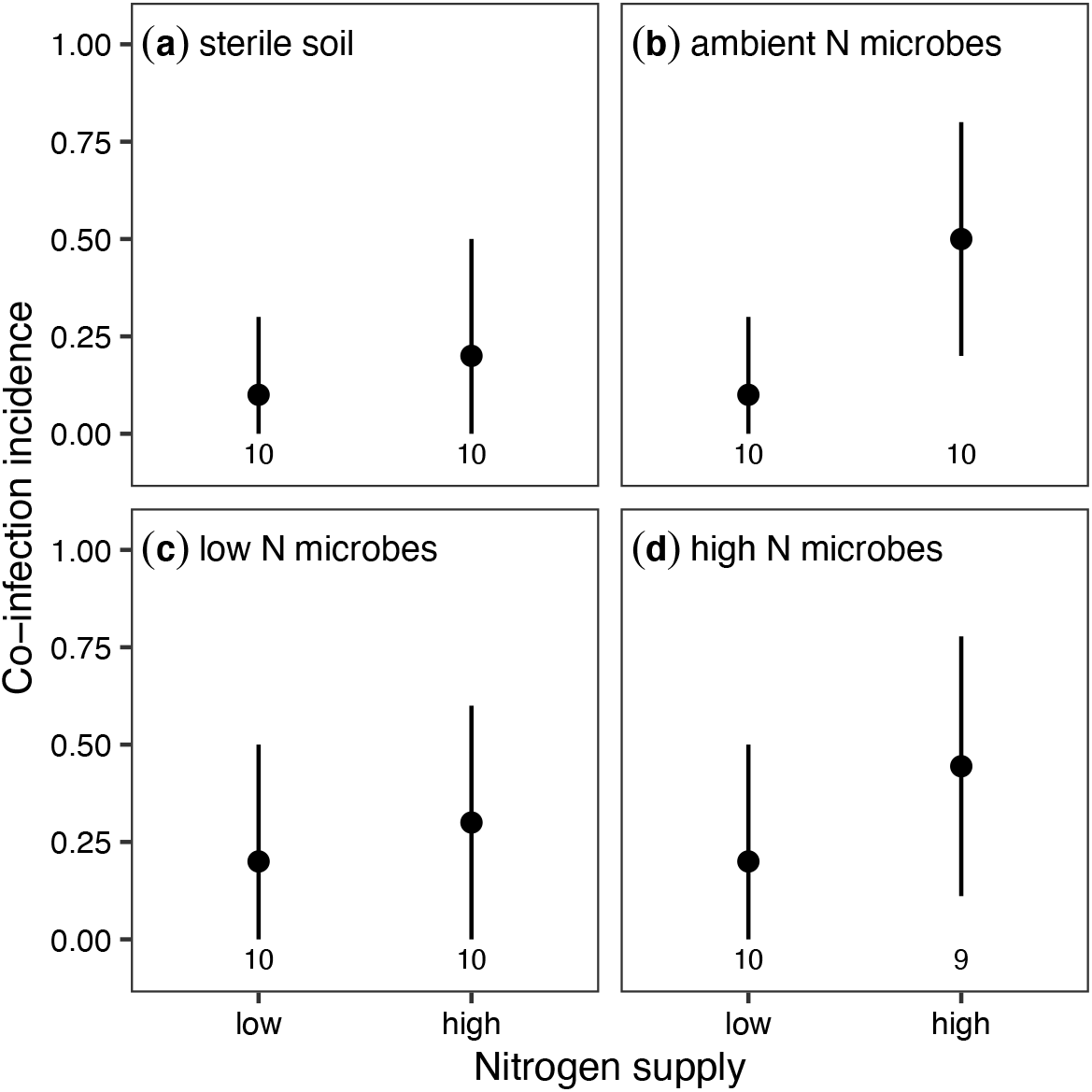
Co-infection incidence (mean ± 95% confidence intervals) of plants grown with (a) sterile soil, (b) microbes exposed to long-term ambient N, (c) microbes exposed to long-term low N, and (d) microbes exposed to long-term high N. Plants were grown with low or high N supply and all were co-inoculated. Corresponding sample sizes are labelled under points and error bars.

### Soil microbes x N and soil microbes x infection: effects on the host

The average aboveground biomass of mock-inoculated plants grown in sterile soil with low N was 0.20 g [0.14 g, 0.28 g] (Fig. **4a**). High N supply increased biomass to 0.37 g, a 90% increase [-0.75%, 185%] (Fig. **2a**,Table **4**). Infection and soil microbes did not significantly affect aboveground biomass (Table **4**). CYDV-RPV infection reduced aboveground biomass on average, but the 95% credible intervals included zero. Only 9.8% of simulated datasets of the same sample sizes used in the experiment had 95% credible intervals that omitted zero. In contrast, sample sizes ten times of those in the experiment produced 95% credible intervals that omitted zero in 72.3% of simulated datasets.

**Table 4.** Linear model summary of log-transformed aboveground biomass (*n* = 154).

| Variable<sup>a</sup> | Estimate | Std. Error | 95% CI |  | Rhat |
| --- | --- | --- | --- | --- | --- |
|  |  |  | Lower | Upper |  |
| **intercept<sup>b</sup>** | **-1.61** | **0.19** | **-1.97** | **-1.24** | **1.00** |
| **N supply** | **0.61** | **0.25** | **0.11** | **1.11** | **1.00** |
| BYDV-PAV infection | 0.19 | 0.26 | -0.31 | 0.69 | 1.00 |
| CYDV-RPV infection | -0.23 | 0.28 | -0.79 | 0.32 | 1.00 |
| soil ambient N | 0.03 | 0.30 | -0.56 | 0.62 | 1.00 |
| soil low N | 0.51 | 0.26 | -0.01 | 1.02 | 1.00 |
| soil high N | 0.09 | 0.27 | -0.45 | 0.62 | 1.00 |
| N supply x BYDV-PAV | -0.13 | 0.41 | -0.93 | 0.66 | 1.00 |
| N supply x CYDV-RPV | -0.06 | 0.37 | -0.79 | 0.68 | 1.00 |
| soil ambient N x N supply | 0.25 | 0.39 | -0.52 | 1.00 | 1.00 |
| soil low N x N supply | -0.22 | 0.37 | -0.94 | 0.51 | 1.00 |
| soil high N x N supply | 0.14 | 0.37 | -0.58 | 0.85 | 1.00 |
| soil ambient N x BYDV-PAV | -0.52 | 0.42 | -1.34 | 0.31 | 1.00 |
| soil low N x BYDV-PAV | -0.41 | 0.41 | -1.22 | 0.42 | 1.00 |
| soil high N x BYDV-PAV | 0.07 | 0.38 | -0.67 | 0.81 | 1.00 |
| soil ambient N x CYDV-RPV | -0.30 | 0.42 | -1.13 | 0.53 | 1.00 |
| soil low N x CYDV-RPV | -0.39 | 0.41 | -1.20 | 0.43 | 1.00 |
| soil high N x CYDV-RPV | 0.12 | 0.41 | -0.68 | 0.93 | 1.00 |
| soil ambient N x N supply x BYDV-PAV | 0.03 | 0.63 | -1.19 | 1.28 | 1.00 |
| soil low N x N supply x BYDV-PAV | -0.19 | 0.59 | -1.35 | 0.95 | 1.00 |
| soil high N x N supply x BYDV-PAV | -0.26 | 0.56 | -1.35 | 0.83 | 1.00 |
| soil ambient N x N supply x CYDV-RPV | 0.19 | 0.58 | -0.93 | 1.34 | 1.00 |
| soil low N x N supply x CYDV-RPV | 0.00 | 0.56 | -1.08 | 1.10 | 1.00 |
| soil high N x N supply x CYDV-RPV | -0.67 | 0.57 | -1.79 | 0.44 | 1.00 |

<sup>a</sup>Variables with quantile-based 95% credible intervals (CI) that omit zero are in bold.
<sup>b</sup>Intercept factor levels: low N supply, mock inoculation, sterile soil

The average leaf chlorophyll content of mock-inoculated plants grown in sterile soil with low N was 23 SPAD [21 SPAD, 25 SPAD] (Fig. **3a**). High N supply increased leaf chlorophyll content to 27 SPAD, a 19% increase [4%, 35%] (Fig. **3a**, Table **5**). Infection did not significantly affect leaf chlorophyll content of plants grown in sterile soil (Fig. **3a**, Table **5**), but BYDV-PAV infection reduced leaf chlorophyll content from 24 SPAD to 19 SPAD, a 21% decrease [-34%, −7.3%], for plants grown with microbes from ambient N field soils (Fig. **3b**). However, when plants were grown with high N supply and ambient N microbes, BYDV-PAV infection did not affect leaf chlorophyll content (estimated change relative to single infection: −7% [-24%, 9%], Fig. **3b**), leading to a significant three-way interaction among ambient N microbes, N supply, and BYDV-PAV infection (Table **5**).

**Table 5.** Linear model summary of log-transformed leaf chlorophyll content (*n* = 154).

| Variable<sup>a</sup> | Estimate | Std. Error | 95% CI |  | Rhat |
| --- | --- | --- | --- | --- | --- |
|  |  |  | Lower | Upper |  |
| **intercept<sup>b</sup>** | **3.13** | **0.05** | **3.04** | **3.23** | **1.00** |
| **N supply** | **0.17** | **0.07** | **0.04** | **0.30** | **1.00** |
| BYDV-PAV infection | 0.08 | 0.07 | -0.05 | 0.21 | 1.00 |
| CYDV-RPV infection | 0.06 | 0.07 | -0.09 | 0.20 | 1.00 |
| soil ambient N | 0.05 | 0.08 | -0.11 | 0.20 | 1.00 |
| soil low N | 0.00 | 0.07 | -0.14 | 0.14 | 1.00 |
| soil high N | 0.02 | 0.07 | -0.13 | 0.16 | 1.00 |
| N supply x BYDV-PAV | -0.20 | 0.11 | -0.41 | 0.01 | 1.00 |
| N supply x CYDV-RPV | -0.15 | 0.10 | -0.34 | 0.05 | 1.00 |
| soil ambient N x N supply | -0.05 | 0.10 | -0.24 | 0.16 | 1.00 |
| soil low N x N supply | 0.04 | 0.09 | -0.14 | 0.23 | 1.00 |
| soil high N x N supply | 0.04 | 0.10 | -0.15 | 0.23 | 1.00 |
| **soil ambient N x BYDV-PAV** | **-0.32** | **0.11** | **-0.53** | **-0.11** | **1.00** |
| soil low N x BYDV-PAV | -0.14 | 0.11 | -0.35 | 0.07 | 1.00 |
| soil high N x BYDV-PAV | -0.06 | 0.10 | -0.25 | 0.14 | 1.00 |
| soil ambient N x CYDV-RPV | -0.11 | 0.11 | -0.32 | 0.11 | 1.00 |
| soil low N x CYDV-RPV | -0.04 | 0.11 | -0.25 | 0.17 | 1.00 |
| soil high N x CYDV-RPV | -0.04 | 0.11 | -0.24 | 0.17 | 1.00 |
| **soil ambient N x N supply x BYDV-PAV** | **0.37** | **0.16** | **0.05** | **0.69** | **1.00** |
| soil low N x N supply x BYDV-PAV | 0.06 | 0.15 | -0.24 | 0.35 | 1.00 |
| soil high N x N supply x BYDV-PAV | 0.03 | 0.15 | -0.25 | 0.32 | 1.00 |
| soil ambient N x N supply x CYDV-RPV | 0.22 | 0.15 | -0.08 | 0.52 | 1.00 |
| soil low N x N supply x CYDV-RPV | 0.11 | 0.14 | -0.17 | 0.39 | 1.00 |
| soil high N x N supply x CYDV-RPV | 0.04 | 0.15 | -0.26 | 0.33 | 1.00 |

<sup>a</sup>Variables with quantile-based 95% credible intervals (CI) that omit zero are in bold.
<sup>b</sup>Intercept factor levels: low N supply, mock inoculation, sterile soil

**Figure 4.**
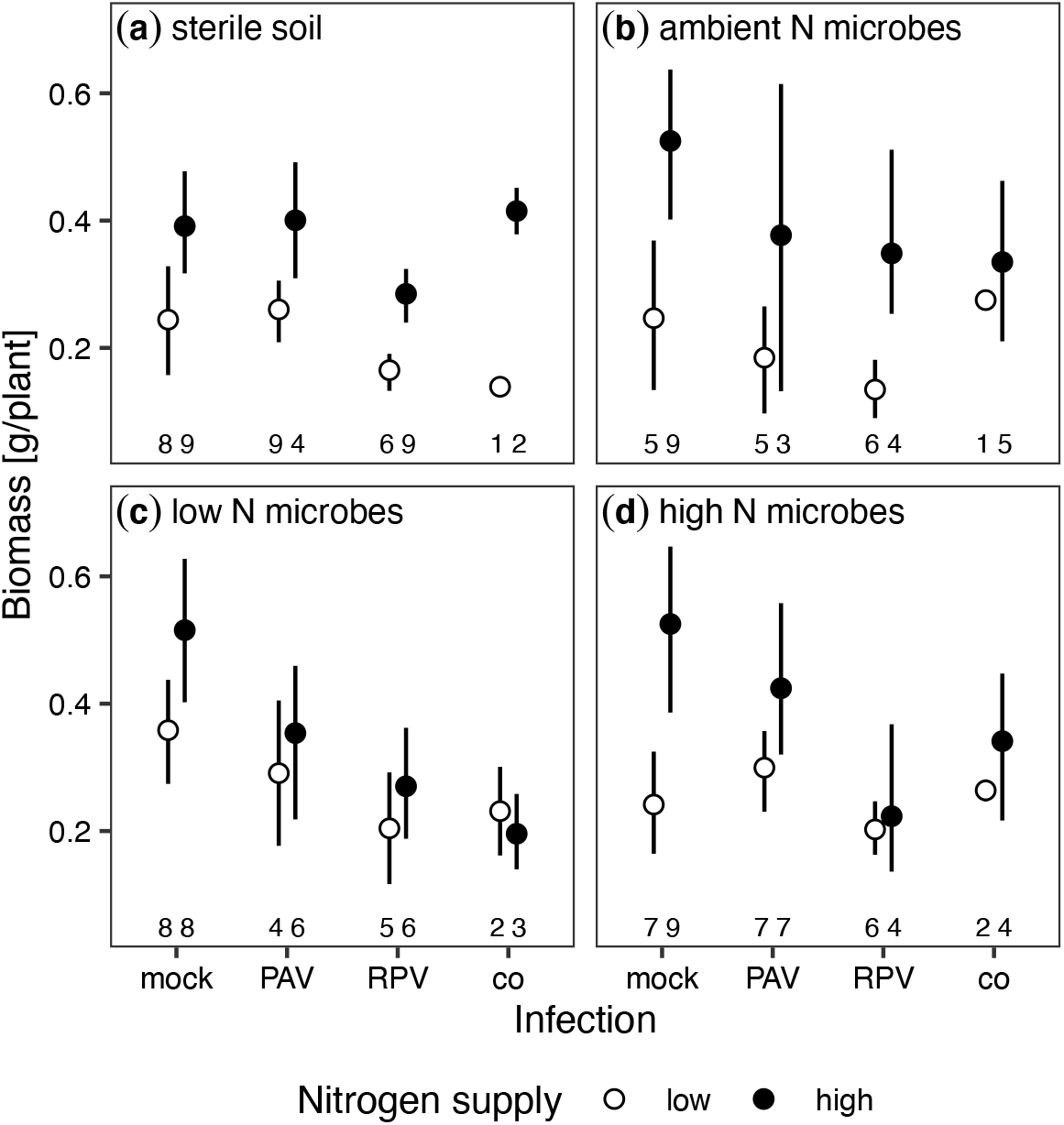
Biomass (g/plant, mean ± 95% confidence intervals) of plants grown with (a) sterile soil, (b) microbes exposed to long-term ambient N, (c) microbes exposed to long-term low N, and (d) microbes exposed to long-term high N. Plants were grown with low or high N supply and infected with mock inoculation, BYDV-PAV, CYDV-RPV, or both (co-infection). Corresponding sample sizes are labelled under points and error bars.

**Figure 5.**
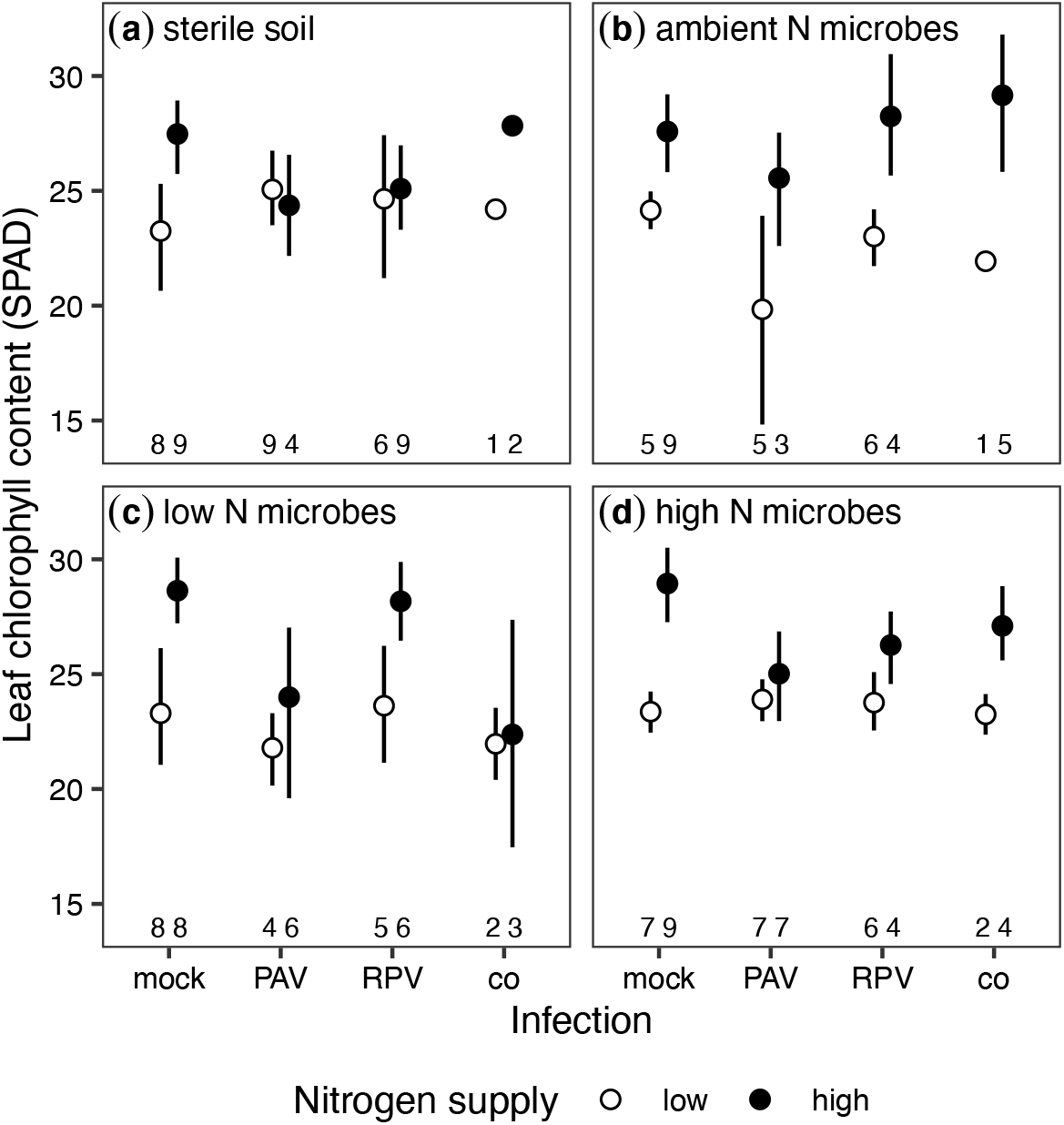
Leaf chlorophyll content (SPAD, mean ± 95% confidence intervals) of plants grown with (a) sterile soil, (b) microbes exposed to long-term ambient N, (c) microbes exposed to long-term low N, and (d) microbes exposed to long-term high N. Plants were grown with low or high N supply and infected with mock inoculation, BYDV-PAV, CYDV-RPV, or both (co-infection). Corresponding sample sizes are labelled under points and error bars.

## Discussion

Human activities have dramatically increased N supply to terrestrial ecosystems, with consequences for plant communities, plant–pathogen interactions, and pathogen–pathogen interactions (Elser *et al*., 2007; Veresoglou *et al*., 2013; Smith, 2014; Midolo *et al*., 2019). Although soil microbes can mediate N availability, soilborne pathogens, and plant defenses (van der Heijden *et al*., 2008; Kuzyakov & Xu, 2013; Mauch-Mani *et al*., 2017), the indirect effects of soil microbes on aboveground pathogen communities and host-pathogen interactions have received little attention. Our experimental manipulation of N supply, soil microbe inoculum, and virus infection demonstrated that soil microbes can influence aboveground plant–pathogen and pathogen–pathogen interactions and mediate the impact of infection on plant traits. Specifically, soil microbes exposed to long-term low N enrichment mediated the effects of N supply and co-inoculation on infection incidence and plant chlorophyll content. Surprisingly, soil microbes did not affect aboveground plant biomass, and they had different impacts on the incidence of two related viral pathogens.

Soil microbes reduced BYDV-PAV incidence and counteracted the negative effects of N supply and co-inoculation on BYDV-PAV incidence. This is the first demonstration that soil microbes can mediate the effects of N supply on a non-soilborne plant pathogen. Of the three general mechanisms through which soil microbes can affect plant–pathogen interactions— modified access to resources (Schimel & Bennett, 2004; van der Heijden *et al*., 2008; Kuzyakov & Xu, 2013), interference with pathogens in the soil (Lugtenberg & Kamilova, 2009; Berendsen *et al*., 2012; Schlatter *et al*., 2017), and induced plant defenses (van Loon *et al*., 1998; Pieterse *et al*., 2014; Mauch-Mani *et al*., 2017)—the third is the most likely to apply to our study. Soil microbes did not significantly affect plant biomass and only affected leaf chlorophyll content in one treatment, suggesting that soil microbes did not generally affect plant access to resources. Because BYDV-PAV and CYDV-RPV are obligately transmitted to plants by aphid vectors (D’Arcy & Burnett, 1995), soil microbes could only affect virus incidence indirectly via effects on the plant, ruling out the second mechanism. Induction of plant defenses could explain why BYDV-PAV incidence was significantly lower when plants were grown with microbes from low N soils. Further, ambient and low N microbes negated the effects of co-inoculation and N supply on BYDV-PAV incidence and the combined effect of co-inoculation and N supply on BYDV-PAV incidence were sub-additive. These interactions are consistent with these factors affecting BYDV-PAV incidence through a common mechanism, such as host defenses.

Furthermore, increased N supply reduced BYDV-PAV incidence in singly inoculated plants grown in sterile soil. Higher N availability may have decreased plant susceptibility to infection, which has been demonstrated for other, usually necrotrophic, pathogens (Dordas, 2009; Vega *et al*., 2015). In a field experiment, N supply decreased BYDV-PAV incidence only when P supply was high, which suggests that the stoichiometry of nutrient supply (e.g., N:P) influences plant–pathogen interactions rather than the absolute supply (Borer *et al*., 2014). Our results may therefore indicate that BYDV-PAV infection was more successful when plants were grown with higher P:N supply, perhaps due to higher within-host virus replication, as has been demonstrated in aquatic systems with P:C stoichiometry (Clasen & Elser, 2007; Frost *et al*., 2008). However, studies that have measured within-host BYDV-PAV titer have not found a positive effect of P or P:N (Rúa *et al*., 2013; Whitaker *et al*., 2015; Lacroix *et al*., 2017; Kendig *et al*., 2020). The mechanism behind reduced BYDV-PAV incidence with higher N supply therefore requires a closer examination of plant defenses, within-host dynamics, and plant–vector interactions. Nitrogen supply no longer reduced infection incidence when plants were co-inoculated. Interestingly, N supply and co-inoculation also interacted to affect virus incidence in a study conducted by Lacroix et. al. (2014), except that the interaction affected CYDV-RPV incidence rather than BYDV-PAV. These results suggest that BYDV-PAV and CYDV-RPV may interact within hosts and that nutrient supply may modify their interactions, analogous to pathogens in animal and human systems (Smith & Holt, 1996; Smith, 2014).

The long-term, field fertilization treatments that shaped the soil microbial communities modified plant–microbe and plant–pathogen interactions. Only soil microbes exposed to multiple decades of low N fertilization affected BYDV-PAV incidence in singly inoculated plants. In addition, co-inoculation only reduced BYDV-PAV incidence in plants grown with sterile soil or microbes from long-term high N fertilization conditions. If soil microbes affected BYDV-PAV incidence through induced plant defenses, these results suggest that the long-term N enrichment could indirectly influence plant defenses via the soil microbial community. Specifically, microbes that have experienced lower long-term N fertilization tended to reduce BYDV-PAV incidence more than microbes that experienced long-term high N fertilization. This result contrasts with previous studies that have demonstrated that higher nutrient availability increased the disease suppressive activity of plant–associated microbes (Wiggins & Kinkel, 2005; Berg & Koskella, 2018). We also found that BYDV-PAV infection only reduced leaf chlorophyll content when plants were grown with microbes exposed to long-term ambient N. This result suggests that not only can the microbial community mediate the success of BYDV-PAV infection, but it also can mediate some of the symptoms of infection experienced by plants. We do not know whether long-term N fertilization shifted the composition or function of the soil microbial communities (Leff *et al*., 2015; Klinger *et al*., 2016; Chen *et al*., 2019), but subsequent studies could characterize microbial taxa and function associated with changes in BYDV-PAV infection.

We selected the host species and virus species because of their importance for agriculture (Mckirdy *et al*., 2002; Riedell *et al*., 2007) and the wide knowledge base provided by previous studies (Carrigan *et al*., 1983; Baltenberger *et al*., 1987; Power *et al*., 1991; Erion & Riedell, 2012; Lacroix *et al*., 2014). However, the host species does not naturally co-occur with the soil microbial communities sampled in this study, which may have limited the observed effects of microbe inoculum on plant growth and plant–pathogen interactions (Essarioui *et al*., 2020). Indeed, studies that have used co-occurring plant species and soil microbial communities have found statistically significant effects of the N fertilization history of microbes on plant biomass (Johnson, 1993; Weese *et al*., 2015). Therefore, our study may have isolated the effects of soil microbes that are generalists or that affect plant–pathogen interactions without requiring co-evolved interactions. A follow-up study may consider exploring the relationship of nutrients, plant pathogens, and soil microbes from communities in which the species naturally occur. In addition to no effect of soil microbes on plant biomass, we also found no effect of infection on plant biomass. BYDV-PAV and CYDV-RPV infections typically reduce plant biomass (Baltenberger *et al*., 1987; Erion & Riedell, 2012), but the effects of plant pathogens are highly dependent on environmental conditions (Barrett *et al*., 2009). Additionally, based on simulated datasets, our results demonstrated that our study’s sample sizes may have impeded our ability to detect small changes in biomass.

Barley and Cereal Yellow Dwarf Viruses are vectored by aphids, which may mediate the effects of soil nutrients and soil microbes on plant–pathogen and pathogen–pathogen interactions. For example, increasing N supply to plants can increase or decrease the length of time that aphids feed (Nowak & Komor, 2010; Bogaert *et al*., 2017), which affects the probability that viruses successfully infect plants (Power *et al*., 1991). Soil microbial communities can also influence the feeding behavior and population dynamics of aphids (Pineda *et al*., 2010). For example, soil microbes can increase or decrease the weight, body size, and intrinsic growth rate of aphids (Hol *et al*., 2010; Pineda *et al*., 2012; Hackett *et al*., 2013). Higher aphid population densities may increase the co-infection incidence of B/CYDVs (Seabloom *et al*., 2009). In addition, N supply and soil microbial communities may interact to affect aphid vectors because the effects of microbes on insect herbivores tend to be stronger when plants experience abiotic stress (Pineda *et al*., 2013). For instance, a previous study demonstrated that other soil-dwelling organisms, nematodes, affected aphid population growth rates and plant preference only when N was limited (Kutyniok *et al*., 2014). To evaluate the role of aphids in pathogen responses to N supply and soil microbial communities, future studies could investigate the effects of N and microbes on aphid feeding duration in the lab and aphid population dynamics in the field.

Our experiment provided a first demonstration that soil microbes can affect the incidence of an insect-vectored virus, BYDV-PAV, in a plant host, *A. sativa*. Because soil microbes mediated the effects of contemporary N supply, and the N fertilization history of soil microbes impacted both incidence and chlorophyll of infected plants, our results suggest that inferences about how N enrichment modifies plant pathogens based on laboratory experiments will depend on the role of soil microbes. Further, our results suggest that high N fertilization could reduce the pathogen suppressive effects of soil microbes for some plant–pathogen pairs, potentially leading to more widespread infection under field conditions with elevated N supply. This work demonstrates the important indirect role of soil microbial communities for infection outcomes, pointing to an exciting new frontier: examining the generality and context-dependence of these results, and the indirect role of soil microbes in the high variation in plant–microbe (Smith & Goodman, 1999) and plant–pathogen (Hoffland *et al*., 2000) interactions.

## Acknowledgements

This work was supported by the NSF program in Ecology and Evolution of Infectious Diseases (grant DEB-1015805). This work also was supported by grants from the US National Science Foundation Long-Term Ecological Research Program (LTER) including DEB-1234162 and DEB-1831944. Further support was provided by the Cedar Creek Ecosystem Science Reserve, the Minnesota Supercomputer Institute, and the University of Minnesota.

## Author Contribution

All authors contributed to the design of the research, CE performed the research and data collection, CAE and AEK performed the data analysis and wrote the first draft of the manuscript, and all authors contributed to manuscript revisions.

## Data Availability

Data will be accessible through the Environmental Data Initiative upon manuscript publication.

