## Supplementary material for "Soil microbes mediate the effects of nitrogen supply and co-inoculation on Barley Yellow Dwarf Virus in *Avena sativa*": Table S1

Easterday *et al.*

**Table S1.** Modified Hoagland solution used to manipulate N supply rate. The low and high concentrations of $NH_4NO_3$ are 0.2% and 10% of half-strength Hoagland solution, respectively.

| Compound | Molar Mass | Concentration (µM) |
|---|---|---|
| $K_2SO_4$ | 174.25 | 1250 |
| $MgSO_4{\cdot}7H_2O$ | 246.48 | 1000 |
| $KH_2PO_4$ | 136.09 | 1 |
| $CaSO_4{\cdot}2H_2O$ | 172.17 | 2000 |
| $NH_4NO_3$ | 80.04 | 7.5 (375 for high N) |
| KCl | 74.56 | 25 |
| $H_3BO_3$ | 61.83 | 12.5 |
| $MnSO_4{\cdot}H_2O$ | 169.02 | 1 |
| $ZnSO_4{\cdot}7H_2O$ | 278.56 | 1 |
| $CuSO_4{\cdot}5H_2O$ | 249.69 | 0.25 |
| $H_2MoO_4{\cdot}(H_2O)$ | 161.95 | 0.25 |
| NaFeEDDHA (6% Fe) | 434.8 | 10 |

**Table S2.** Plants with virus infections inconsistent with their inoculation treatment. Total sample sizes are in parentheses.

| Inoculation treatment | BYDV-PAV | CYDV-RPV |
|---|---|---|
| Mock (77) | 2 | 12 |
| CYDV-RPV (78) | 3 | |
| BYDV-PAV (74) | | 14 |

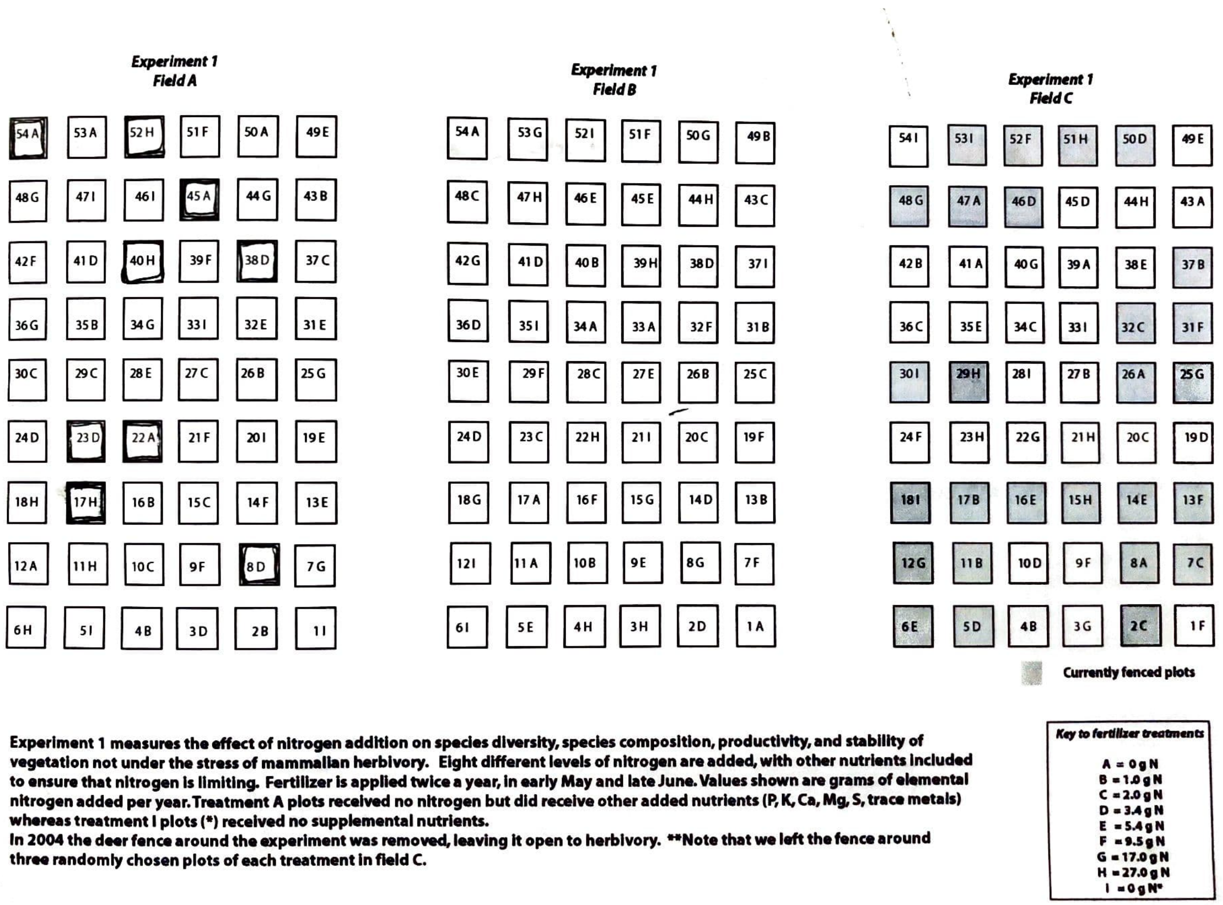


**Fig. S1.** Map of plots within Cedar Creek Ecosystem Reserve experiment where soil cores were sampled. Soil was collected from Field A in each of the plots with heavy borders (54A, 52H, 45A, 40H, 38D, 23D, 22A, 17H, and 8D). Plots were selected using a random number generator. To select locations within a plot where soil cores were extracted, we face away from the plot and threw a ball over our shoulder.
